# VUStruct: a compute pipeline for high throughput and personalized structural biology

**DOI:** 10.1101/2024.08.06.606224

**Authors:** Christopher W. Moth, Jonathan H. Sheehan, Abdullah Al Mamun, R. Michael Sivley, Alican Gulsevin, David Rinker, Undiagnosed Diseases Network, John A. Capra, Jens Meiler

**Affiliations:** Departments of Chemistry, Pharmacology, and Biomedical Informatics; Center for Structural Biology and Institute of Chemical Biology; Vanderbilt Univ., Nashville, TN 37232, USA; Division of Infection Diseases, Milliken Dept. of Internal Medicine, Washington Univ. of Medicine in St. Louis, MO 63110, USA; Biomedical Informatics at 5Prime Sciences, Montreal, Canada; Department of Pharmaceutical Sciences, College of Pharmacy and Health Sciences, Butler University, Indianapolis, IN 46208, USA; Department of Biological Sciences, Evolutionary Studies Initiative; Vanderbilt Univ., Nashville, TN 37232, USA; Bakar Computational Health Science Institute and Department of Epidemiology and Biostatistics, Univ. of California San Francisco, CA 94143, USA; Leipzig University Medical School, Institute for Drug Discovery, Brüderstraße 34, 04103 Leipzig, Germany

## Abstract

Effective diagnosis and treatment of rare genetic disorders requires the interpretation of a patient’s genetic variants of unknown significance (VUSs). Today, clinical decision-making is primarily guided by gene-phenotype association databases and DNA-based scoring methods. Our web-accessible variant analysis pipeline, VUStruct, supplements these established approaches by deeply analyzing the downstream molecular impact of variation in context of 3D protein structure. VUStruct’s growing impact is fueled by the co-proliferation of protein 3D structural models, gene sequencing, compute power, and artificial intelligence.

Contextualizing VUSs in protein 3D structural models also illuminates longitudinal genomics studies and biochemical bench research focused on VUS, and we created VUStruct for clinicians and researchers alike. We now introduce VUStruct to the broad scientific community as a mature, web-facing, extensible, High-Performance Computing (HPC) software pipeline.

VUStruct maps missense variants onto automatically selected protein structures and launches a broad range of analyses. These include energy-based assessments of protein folding and stability, pathogenicity prediction through spatial clustering analysis, and machine learning (ML) predictors of binding surface disruptions and nearby post-translational modification sites. The pipeline also considers the entire input set of VUS and identifies genes potentially involved in digenic disease.

VUStruct’s utility in clinical rare disease genome interpretation has been demonstrated through its analysis of over 175 Undiagnosed Disease Network (UDN) Patient cases. VUStruct-leveraged hypotheses have often informed clinicians in their consideration of additional patient testing, and we report here details from two cases where VUStruct was key to their solution. We also note successes with academic research collaborators, for whom VUStruct has informed research directions in both computational genomics and wet lab studies.

## Introduction

Clinical diagnosis of the genetic causes of rare diseases is primarily guided by databases of known gene-phenotype associations(1) and computational methods for quantifying the effects of genetic variants. Examples of these methods include GERP, which analyzes evolutionary constraint(2,3); SIFT(4) which performs protein sequence homology analysis; and Polyphen(5), which is additionally trained on observed and predicted protein 3D structural features. While variant effect prediction algorithms have demonstrated utility in distinguishing known pathogenic variants from benign variants across large variant sets, these algorithms suffer from low specificity. Thus, computational methods are often of limited utility for the small sets of pre-filtered variants(6) that are typically analyzed in clinical cases and other applications involving small sets of variants (bench studies of proteins and metabolic pathways, deep mutational scans, etc.)(7). The recent development of more sophisticated ML techniques and larger training data sets has increased the predictive accuracy of scoring algorithms(8,9). Nonetheless, even AlphaMissense’s scores lack reliability in cases of specific variants(8) and can exhibit high false positive rates(9). I.e., with such a high false positive rate, the touted longitudinal statistical significance of these algorithms cannot diagnose an individual patient’s disease, nor reliably identify disruption points in a single protein or metabolic pathway.

Compounding the above caveats, computational variant effect prediction approaches reveal neither molecular nor biological mechanistic hypotheses. Instead, these tools are focused on the broad classification of mutations into pathogenic or benign, a vague partitioning with limited clinical utility. The critical biology of life unfolds in 3D space and time. Yet, scores compress this complex biology into a single number which obscures functional consequences of VUSs and their mechanisms of disease progression.

Mechanistically, variants in protein coding regions can disrupt protein function and cause diseases in various ways. As examples, amino acid substitutions can compromise the subtle energetics of protein folding and thermodynamic stability. Protein-protein interactions can be disrupted, post-translational modifications can be impeded, and metabolic networks can be broken(10).

Recently, computational protein structural analyses have demonstrated the power of mechanistic modeling of variants’ effects to reveal causes of rare disease. For example, structural modeling suggested that a de-novo VUS in KCNC2 (V469L) could block the ion channel pore, impacting the stability of the protein(11). This provided a rational and foundational hypothesis for the mechanism by which V469L causes developmental and epileptic encephalopathies (DEE) symptoms. Structure-based calculations also revealed that a missense variant in MSH2 could destabilize the protein, leading to cellular protein degradation and Lynch-syndrome disorder(12). In these cases, structure-based calculations outperformed the traditionally used genetic disease predictors. The success of the MSH2 study, as well as numerous other single-gene focused analyses, informed the creation of a generalized structure-based workflow for variant classification in the clinic(13). While this workflow provides guidance for 3D structural model selection and curation, the prescribed processes require significant human input. Once selected, structures must be manually forwarded to various external webservers which perform specific calculations, the results of which must still be integrated into final reports and explored with external visualization tools.

We created VUStruct considering these successes and analytical challenges. We hypothesized that an automated pipeline of structure-based calculations could reveal clues of variant structural and functional impacts which, in turn, could lead to the plausible identification of the root causes of rare genetic disorders in many patient cases.

The goal of VUStruct is to provide robust context on the effects of a VUS on protein structure and function - context that enables the development of mechanistic clinical hypotheses about the causes of disease. VUStruct’s automated contextualization of VUSs in protein 3D structural models can also illuminate longitudinal genomics studies and biochemical bench research focused on VUS. The pipeline automatically selects structures, integrates a broad spectrum of established computational approaches, and caps the calculation with holistic case-wide reporting. In contrast to other webservers which display variants on protein structures alongside precomputed and pre-aggregated scores(14,15), VUStruct performs fresh calculations based on queries to current genomic and protein model databases. Many servers require upload of a (single) protein structure file(16) or default to Alphafold-2 models(17) covering only Uniprot(18) canonical transcripts. VUStruct expands the scope of previous methods by integrating analyses of multiple 3D structures per protein and non-canonical transcripts when available. The final computed product is a website that enables drilling-down from a top-level case report, to each transcript, to 3D structure visualization. For each 3D structure, NGLviewer(19) sessions afford not only 3D manipulation of a variant’s spatial environment, but also visualization of the proximity of known pathogenic and benign variants and evolutionary constraint within and between species (PathProx (20,21), ConSurf(22,23) and COSMIS(24)). VUStruct also investigates the potential for combinations of the VUSs to cause disease with DiGePred(25) and DIEP(26) ML algorithms trained to detect digenic disease.

Taken together, these automated and parallel calculations inform the clinic or laboratory at a 3D structural and molecular mechanistic level. VUStruct provides a compelling supplement to the insights gained from conventional genome-based scoring analysis alone, and we report the pipeline’s contribution to two UDN(27,28) patient cases.

### Design and Implementation

The VUStruct computational pipeline is primarily implemented as Python codes which query and filter a wide range of pre-downloaded databases. Additional code launches and monitors the calculations which run inside Singularity(29) containers.

Conceptually, the pipeline runs in five discrete phases following upload of a variant set via the initial web form:

1) Optional pre-processing of human genomic coordinates
2) 3D structure selection and compute job planning
3) Launch of job arrays (for each variant) on the HPC
4) Progress monitoring
5) Report generation

Except for progress monitoring, these phases are depicted in Figure 1 and detailed below.

**Figure 1.**
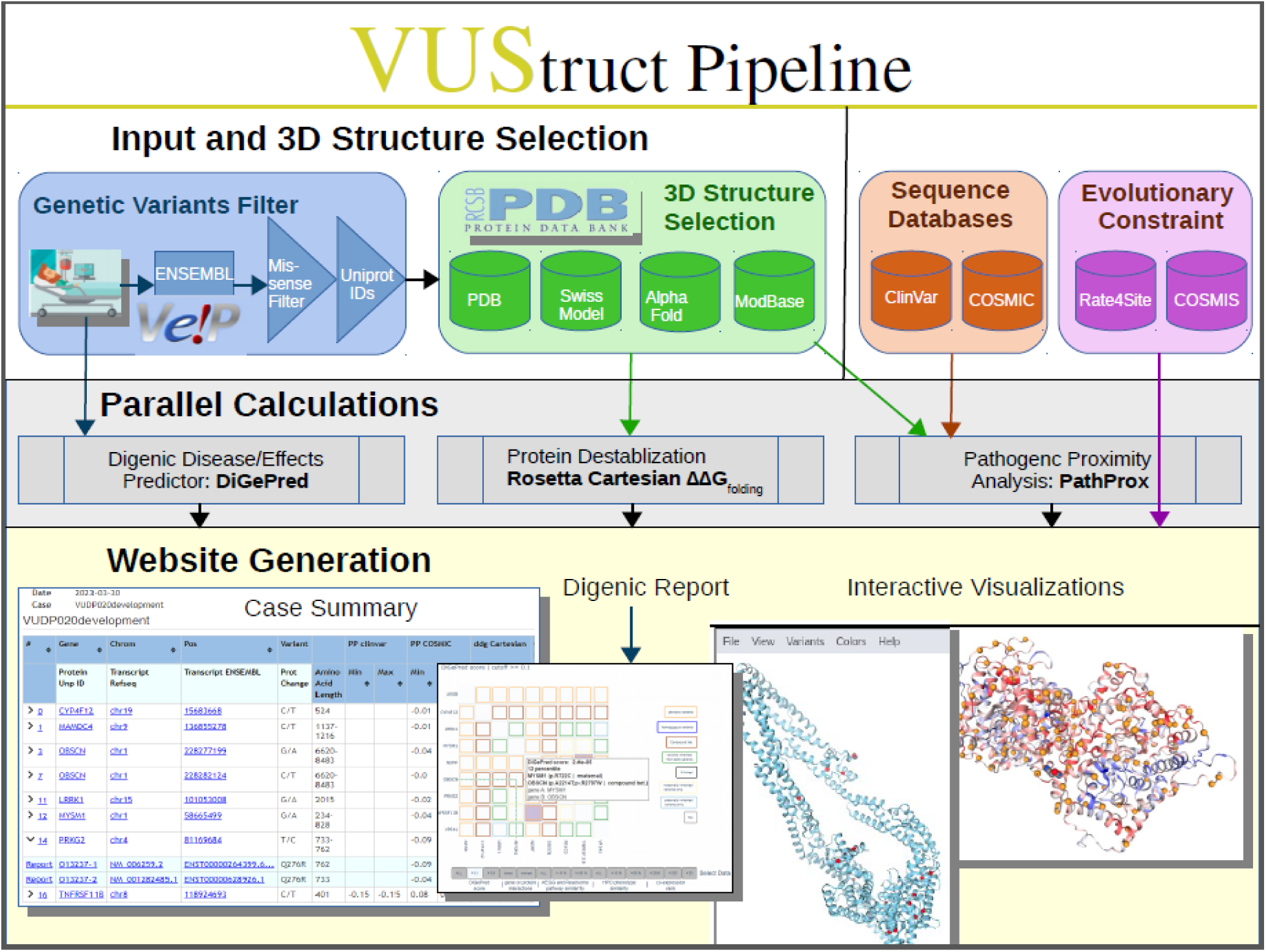
Starting from user-provided variant genomic coordinates (top left), VUStruct identifies missense variants and maps them onto protein structures which VUStruct automatically curates from experimental depositions and model databases. Various parallel calculations are then launched on the HPC, as enumerated in the text.

## 1 Variant Upload and (optional) Genomic Preprocessing

VUStruct supports several input formats, which are first converted into a “vustruct.csv” pipeline-ready, comma-delimited flat file. This “VUStruct CSV” file contains gene names, transcript identifiers, amino acid variants, and parental inheritance when known. When users already know precise transcript identifiers and amino acid changes of interest, then “VUStruct CSV” format may be selected from the outset, and the pipeline will proceed directly to phase 2.

When starting from patient genetic variants (the underlying changes to alleles at chromosome positions) VUStruct converts these data to proteomic impacts. For variants loaded in VCF(30) format, parsed genomic coordinates are fed through the ENSEMBL(31) Variant Effect Predictor (VEP)(32) and missense variants are retained from the VEP output for subsequent structure selection and calculation scheduling. Since the VEP often reports impacts to many predicted transcripts which lack experimental validation, VUstruct restricts its calculations to the subset of returned genomic transcript IDs which cross-reference to Swiss-Prot curated Uniprot(18) protein IDs. Non-canonical transcripts are increasingly found *in-vivo* as proteomics methods evolve(33), and VUstruct includes all of a gene’s curated splice variants. I.e., we incorporate the curated but non-canonical sequences identified in Uniprot with additional “-N” suffixed identifiers.

A challenge in our field is that annotations to the human reference genome(34) are not static. There are relatively frequent amino acid sequence discrepancies between ENSEMBL transcript records and Swiss- Prot curated Uniprot sequences (as cross-referenced by UniParc identifiers in Uniprot’s “Id Mapping” resources). As a practical example, in early 2023, while 48,308 ENSEMBL transcripts cross-referenced to Uniprot sequences perfectly, we also found that 8,500 curated Uniprot IDs had no cross references to any GRCh38 Ensembl Transcript identifiers. 1,166 transcripts had cross-references and the transcript lengths were the same in both databases. However, the amino acid sequences were different. 370 transcripts had varying transcript lengths between between ENSEMBL and Uniprot. Uniprot is constantly working to improve cross-references, and the MANE(35) collaboration is also informing the field. Today, for variants that cannot be immediately processed due to these disconnects, the pipeline reports these problems to the user and the pipeline stops. This provides users with the opportunity to either rework the genomic coordinates input data, or manually download and patch the preprocessor-generated vustruct.csv file for input to phase 2.

## 2 Structure Selection and Compute Job planning

From the Uniprot IDs in the “vustruct.csv” file, the pipeline “plans” the set of calculations by gathering structural information for target proteins. Available experimental structures are mined from the PDB and aligned to current transcripts via the SIFTS database(36,37). SwissModel and ModBase models are integrated (38,39). For mutations on canonical transcripts, AlphaFold(40) models are added to the set of representative structure. The final structure selections minimize redundancy and maximize diversity of experimental techniques, variant-coverage, model confidence and experimental quality metrics. Multimeric complexes are also prioritized in this process.

Below the single directory for the user-provided Case ID, a subdirectory is created for each variant. For each retained structure, calculations are planned, and command line parameters are set for each job. These details are recorded in the workplan.csv of the variant subdirectory. Importantly, planning is entirely independent of the HPC architecture. To ensure that no job conflicts with any other, each user-input Case ID is appended to a Globally Unique Identifier (GUID)(41) and assigned a work directory in the hierarchy of VUStruct/CaseID_and GUID/Transcript ID/3D Structure Type and Id/Calculation Type/Work Directory/. A sibling /Status Directory/ is used by each running job in VUStruct to uniformly communicate progress, competition, or failure to the VUStruct monitor application (described under phase 4).

## 3 Job Launch

To launch the hundred(s) of jobs typically planned for a set of VUS (e.g., from a UDN case or from list of variants from genome sequencing), the pipeline writes submission scripts for the supported cluster environments on the back end (either SLURM(42) or IBM LSF™). Each launched job runs out of a Singularity(29) container. From the container, the bound filesystem of the HPC environment is accessible but the application is otherwise blind to the surrounding HPC API. A single short script, external to the container architecture, launches all the HPC jobs, and records assigned job numbers for downstream monitoring.

The currently launched calculations include:

1) Rosetta ΔΔG_folding_(43–45) estimates the energetic impact of each amino acid substitution on the free energy of protein folding. These two-part calculations are stored in a repository to avoid redundant “relax” steps and save compute time.
2) PathProx (20,21) predicts pathogenic variants when they better fit with clusters of “known pathogenic” sites (mined from Clinvar) (46) vs. randomly placed vs. benign variant sites found in Gnomad (47).
3) ScanNet (48)estimates the likelihood that a variant to disrupt a protein-protein interaction, via an ML algorithm.
4) MusiteDeep (49) predicts protein post-translational (PTM) site modification through a deep-learning framework.
5) Digenic disease interactions are predicted with DigePred(25) and DIEP(26).

Additional suggestions for the interpretation of these outputs are provided in the Supplemental Information.

## 4 Job Monitoring

Over the course of a VUStruct run, the case report is refreshed at 30 minute intervals, to reflect the latest calculated data. The stdout, stderr, and .log files for each individual job are also updated.

The pipeline also informs the user of both overall and individual job progress on the cluster. In a large shared HPC environment, launched jobs are assigned unique job numbers, but do not immediately run. Traditionally, HPC users monitor job progress with a suite of HPC-provided command line tools. Through its web interface, VUStruct interfaces to these tools on the back end, and dynamically reports on job prioritization, submission delays, remaining run time, and resource allocation. These technical status updates are presented to the user via a JavaScript monitor running in the case landing page. This page receives updates from an HPC node via middleware on the web server host.

## 5 Reporting

The pipeline generates a case-wide report as a landing page that combines calculated results for each transcript. As shown in Figure 2, the report also integrates queried scores from AlphaMissense(50), ConSurf(22,23) and COSMIS(24) for all the individual variants. This is followed by digenic analysis outputs.

**Figure 2.**
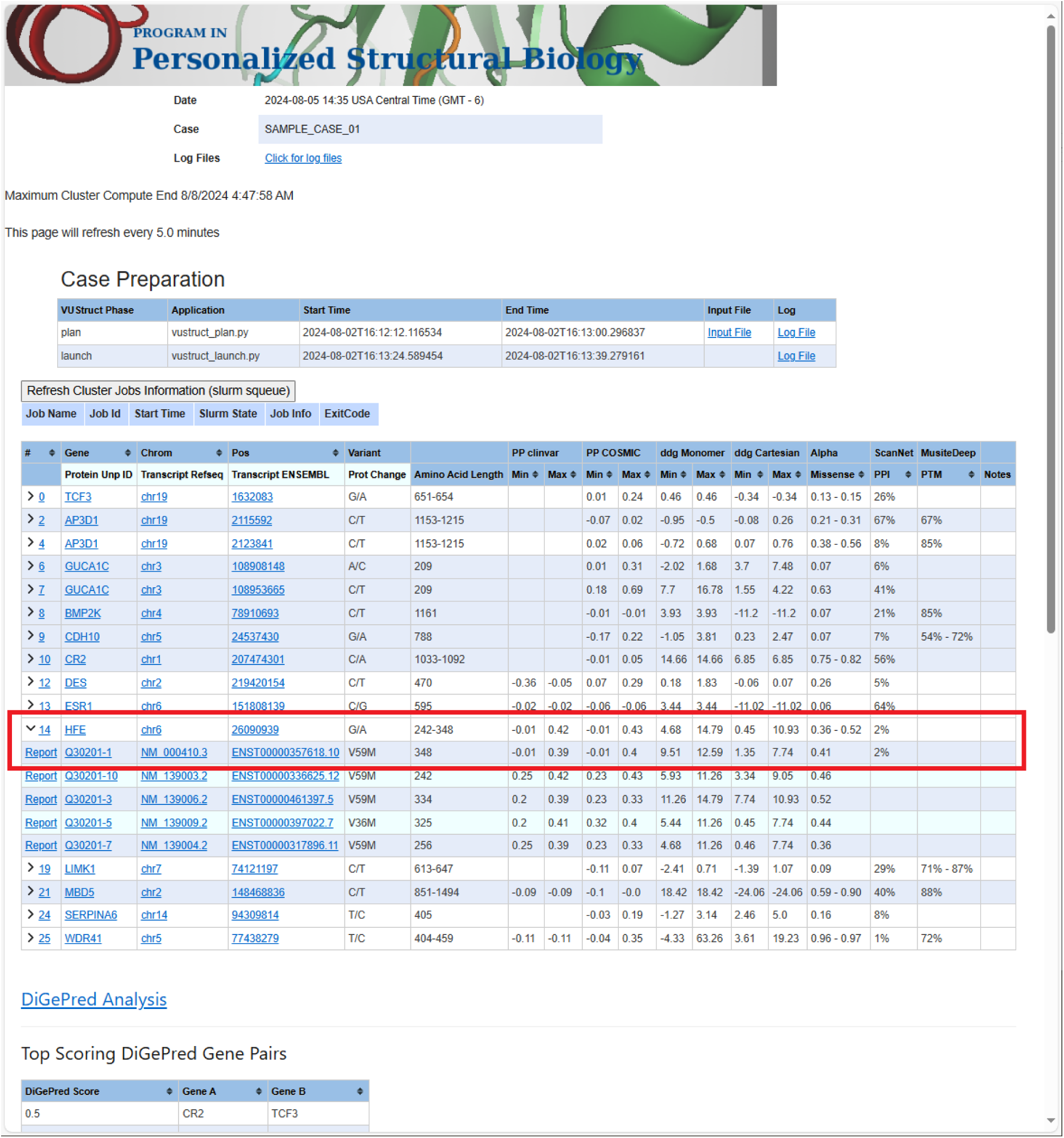
The case report landing page contains a table, in which each row summarizes the range of calculated values for each variant. Ranges arise in some calculations because multiple structures are considered for each transcript. In the case of input genomic coordinates, multiple transcript isoforms are often impacted for each variant. The "Refresh Cluster Jobs Information" box allows detailed monitoring and troubleshooting. The drawn red box shows how summary row 14 (a change to gene HFE on Chrom 6) has been expanded to display five rows for different impacted transcripts. The first of row corresponds to the canonical UniProt isoform. Clicking the "Report" link for that line will display detailed calculations for this variant in the context of that transcript (see next figure).

From the case-wide report, the user may click into specific transcript reports (Figure 3). Clicking into a transcript report presents the user with a PFAM domain graphic(51) followed by a tabular summary of calculation results for the associated structures that were selected in step 2. The “navbar” at top left allows the user to hop to individual 3D structures, where NGL WebViewer(19) sessions are available to inspect the atomic environment of variants (Figure 4). The customized viewer also allows backbone coloring of the various calculated constraint scores, and model confidence.

**Figure 3.**
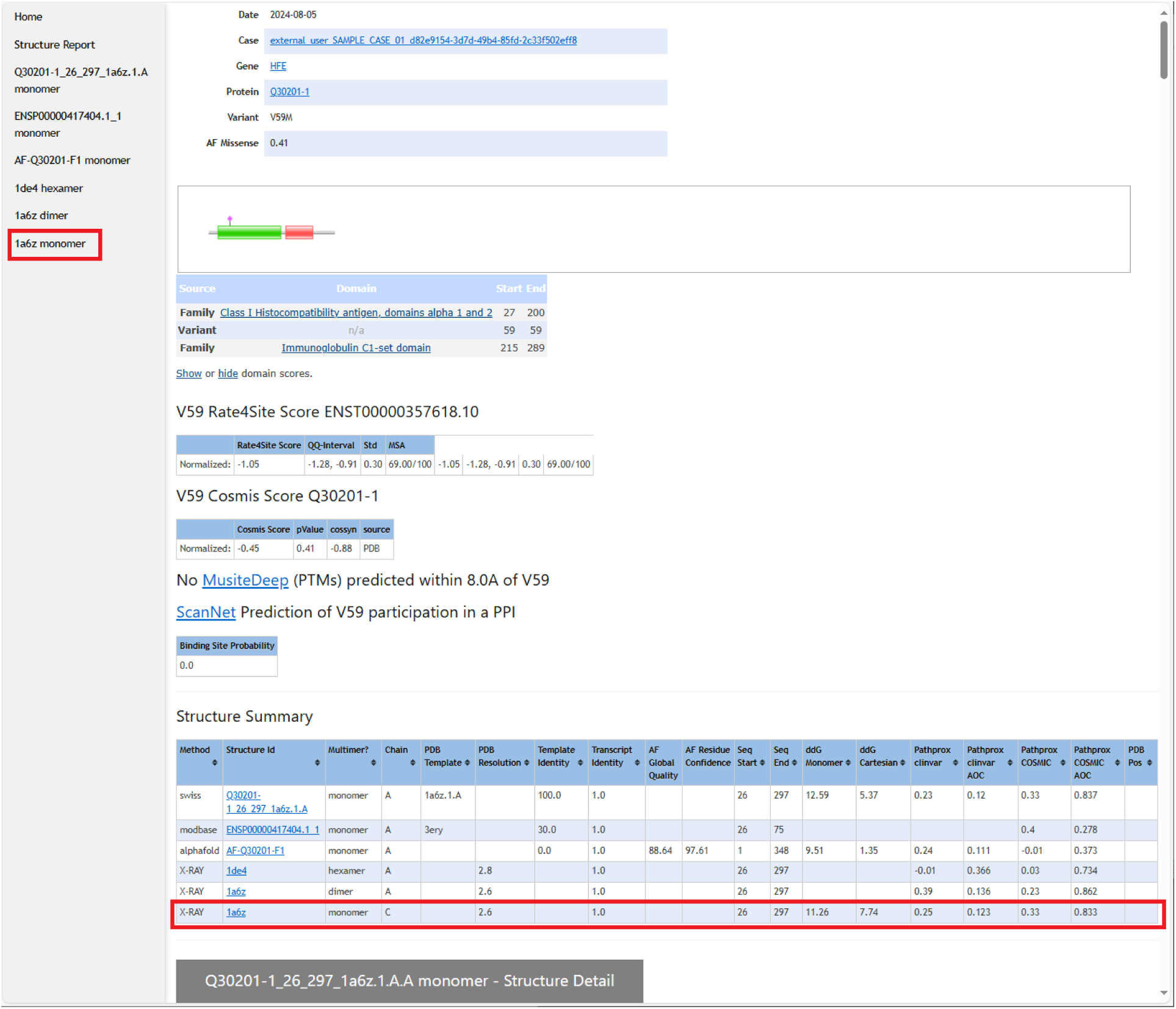
The top of each transcript variant report shows the variant location as a pink diamond in the context of the protein’s PFAM domain(51) annotation. This is followed by results for Rate4Site, COSMIS, MusiteDeep, and ScanNet calculations. A key table is the Structure Summary, which lists all of the structures (from the PDB, MODBASE, SWISS-MODEL database, and AlphaFold database) on which calculations were performed. For example, the highlighted row summarizes all the calculations performed on X-Ray crystal structure 1a6b.pdb, and the highlighted shortcut in the left column leads to the section of the page detailing those results (see next figure).

**Figure 4.**
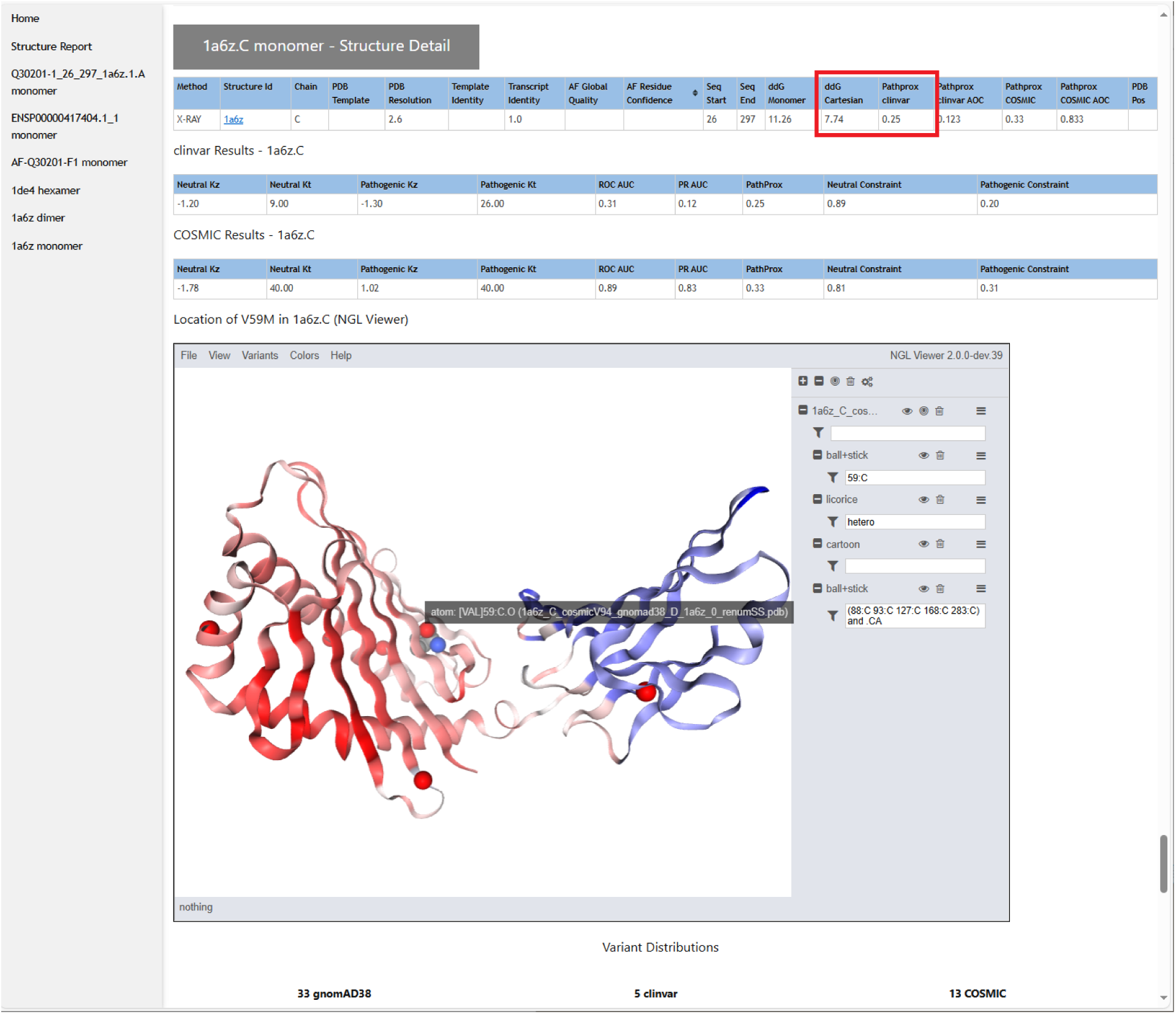
The section of the results page devoted to each structural model shows the results of the ΔΔG and PathProx calculations, highlighted in red, along with associated statistics to judge reliability. Each structure is displayed in a customized and interactive NGL Viewer(19) session. This view can be used to understand the structural context of the variant (highlighted here with a gray text pop-up. For example, one can display all pathogenic and likely pathogenic ClinVar variants (shown as red spheres), or color the model according to AlphaFold confidence or by PathProx score (from low-blue, to high - red - as shown above). Figures to illustrate a structural mechanistic hypothesis can be generated quickly from these images.

The downstream audience for VUStruct case reports is broader than the structural biologists trained to interpret the pipeline’s detailed outputs. Typically, that final audience includes clinicians and geneticists who are primarily interested in whether VUStruct identifies a candidate gene for ongoing consideration, and how pipeline outputs, at high level, inform that recommendation. To communicate the high-level findings of VUStruct succinctly, VUStruct drafts a case summary spreadsheet (Supplemental Information Figure 1). The Supplemental Information also suggests approaches to communicating with clinical partners and includes advice on calculation interpretation.

### Dependencies

VUStruct integrates several externally sourced databases. So that the pipeline can run responsively, and avoid vulnerability to external outages, the supporting databases are locally downloaded, installed, and maintained. The two support pillars of VUStruct are the ENSEMBL GRCh38 PERL API and the UniProt id mapping file. We locally import ENSEMBL’s SQL database, and additionally load UniProt(18) cross- references into SQL tables to speed sequence cross-references between genome and proteome. BASH scripts are additionally provided to aid download of Clinvar(46), COSMIC(52), and gnomAD(47) databases which are mined for PathProx’s mathematical spatial analysis and for web-based visualizations. Several of our predictive calculations integrate sequence constraint, gleaned from both multi-species sequence alignments(22,23) and human population sequences(24). These calculations, along with AlphaMissense(50) predictions, are downloaded as transcriptome-wide precomputations, and are integrated into final reports without the need for cluster launches.

Cited calculations are deployed inside Singularity Containers. Deployment of Rosetta ΔΔG_folding_ (43–45) Cartesian and Monomer calculations requires a free academic or paid commercial license from rosettacommons.org.

### Results/Application of VUStruct in the interpretation of clinical data

We have demonstrated the VUStruct pipeline’s utility in the interpretation of genetic VUS in collaboration with colleagues from the Vanderbilt UDN. The containerized VUStruct software pipeline has been applied to over 150 UDN Vanderbilt UDN patients and 25 Washington University patients. The pipeline provides researchers and clinician geneticists with insights into candidate missense variants in the context of 3D protein atomic structure. In contrast to the many algorithms and websites that perform a single calculation on a single protein variant on a single protein structure, VUStruct is holistic and automated. Our pipeline analyzes a set of patient genetic VUSs and unifies the results under a case-wide report page. VUStruct is also noteworthy for its principled selection of appropriate structures among the growing wealth of available experimental and computational structural models, automated calculation setup and launch, and progress monitoring.

As one illustration of VUStruct’s potential to aid hypothesis generation, we highlight a patient with PASNA syndrome caused by a heterozygous variant in the CACNA1D gene that encodes a Human L-type voltage-gated calcium channel (Cav). Several candidate variants were selected from the patient genome sequencing (GS) data based on phenotype analysis. These variants were submitted to the pipeline and the 3D structure of the corresponding protein was analyzed by different computational methods including Rosetta ΔΔG (43–45), protein-protein interaction (PPI), post-translational modifications (PTM) and digenic predictions (DiGePred) analysis. VUStruct reported that the F767L variant in CACNA1D results in structural destabilization as evidenced by ΔΔG score in Rosetta. Starting from the VUStruct report, we hypothesized that the variant may contribute to the PASNA syndrome and conducted additional Rosetta simulations on the Cav structural model. In follow-up, two different variants F767L and F767S for Cav were used to calculate the ΔΔG in Rosetta using closed state conformation (PDB id: 7UHG (53)). F767S is a known pathogenic variant that causes a gain of function mechanism, and it was used as a positive control for this study. The higher calculated ΔΔG of F767L (∼5.3 Rosetta Energy Units) vs F767S (∼3.9 R.E.U.) suggested that F767L could contribute to at least as much structural disruption as known pathogenic variant F767S for the closed state conformation. Thus we hypothesized that these variants destabilize the closed state, and push conformational equilibrium towards the channel opening state. The search for this crucial finding began with VUStruct analysis and led to the further confirmative analysis to diagnose the possible cause of the PASNA syndrome (54).

A second demonstration of VUStruct’s utility was aiding a diagnosis of Diamond Blackfin anemia (DBA) in a case which could not be explained by simple Mendelian inheritance. The VUStruct report suggested that a missense variant in the RPS19 gene results in a slight stabilization, based on Rosetta ΔΔG. In addition, the proband carried another variant in the RPL27 gene, which DiGePred(25) and DIEP(26) analysis predicted to have a strong digenic interaction with RPS19. These clues helped to focus further structural analysis. We investigated different 80S ribosome structures available in the protein data bank. Although RPS19 and RPL27 are on opposite sides of the complex. It is plausible that T55M in RPS19 changes allosteric interactions between the two proteins, disrupting the 80S ribosome function. These structural analyses inspired further co-segregation and RNA sequencing analysis of the proband. Further analysis of these suggested the proband’s DBA is caused by the digenic interactions between RPS19 and RPL27(55).

### Availability and Future Directions

The website is made available to all, without condition. For those wishing to setup their own pipeline environment, all our code and containers (with one exception), are licensed under the MIT License and can be downloaded from https://github.com/meilerlab/VUStruct. The one exception is the Rosetta ΔΔG module containers, which require Rosetta Commons licensing, available at no charge to academic users.

VUStruct development is continuously fueled by ongoing explosions in available protein 3D structures, genome sequencing, computer power, and artificial intelligence. We are committed to the pipeline’s flexibility and continuous improvement.

One current pipeline limitation is that all calculations are based on sets of single structural models, and the implications of dynamics are not presently considered. Multi-conformer generation is an active area of research(56). We plan to integrate that work into the pipeline, so that more conformational states are sampled. Additionally, we are integrating AlphaFold models for non-canonical transcript sequences(57). Pending its public opening, we hope to mine the AlphaFold 3 repository for its updated structural coverage that includes multimeric complexes(58). Predictions of digenic interactions should benefit from model retraining, given the emergence of new ground truth data sets(59).

## Support and Thanks

This work leveraged the resources provided by the Vanderbilt Advanced Computing Center for Research and Education (ACCRE), a collaboratory operated by and for Vanderbilt faculty. ACCRE is comprised of over 3,000 researchers from more than 40 campus departments.

The pipeline would not have been possible without the energetic and helpful support from staff at Uniprot, ENSEMBL, SwissModel, and Modbase.

## UDN Consortium

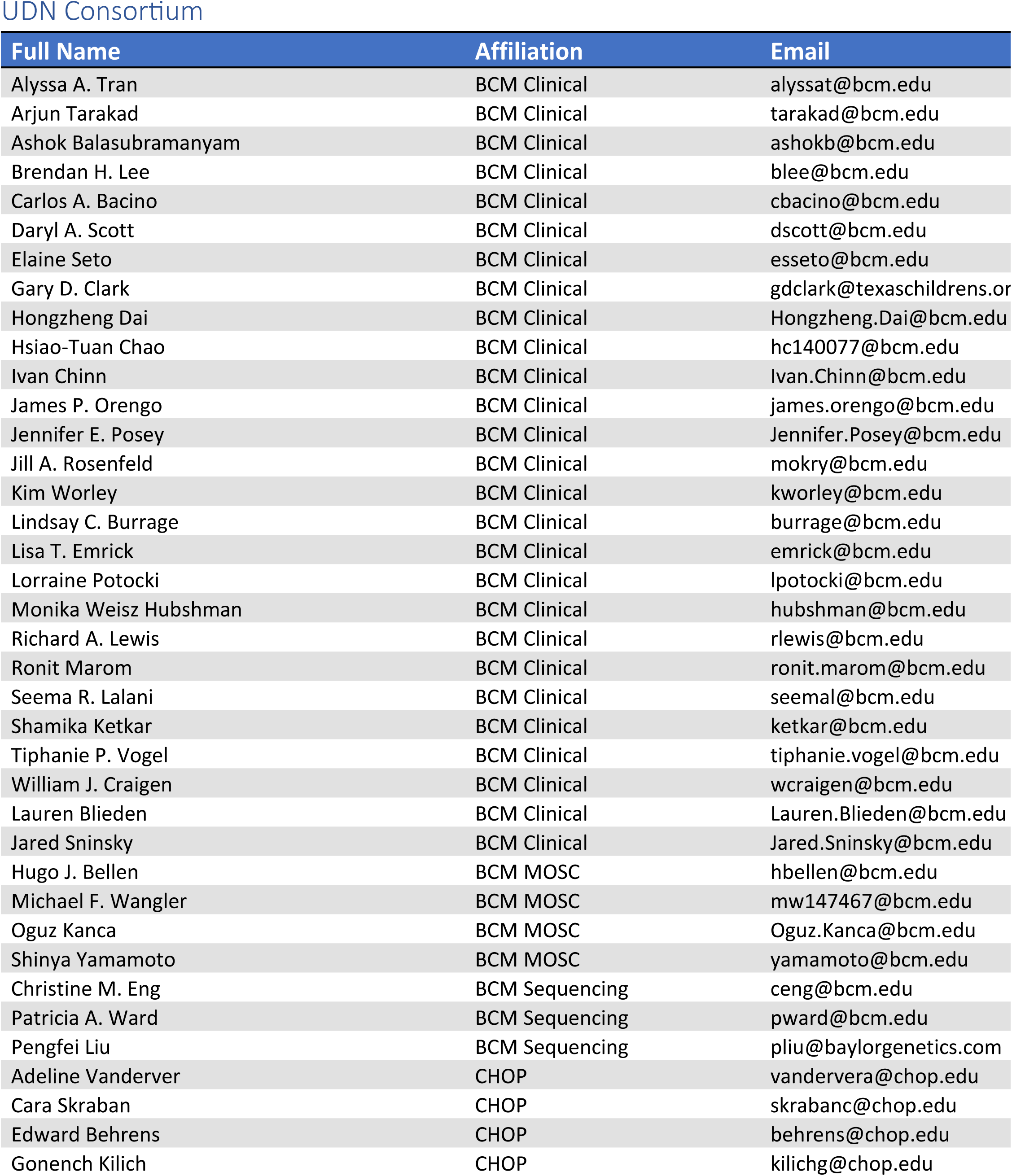
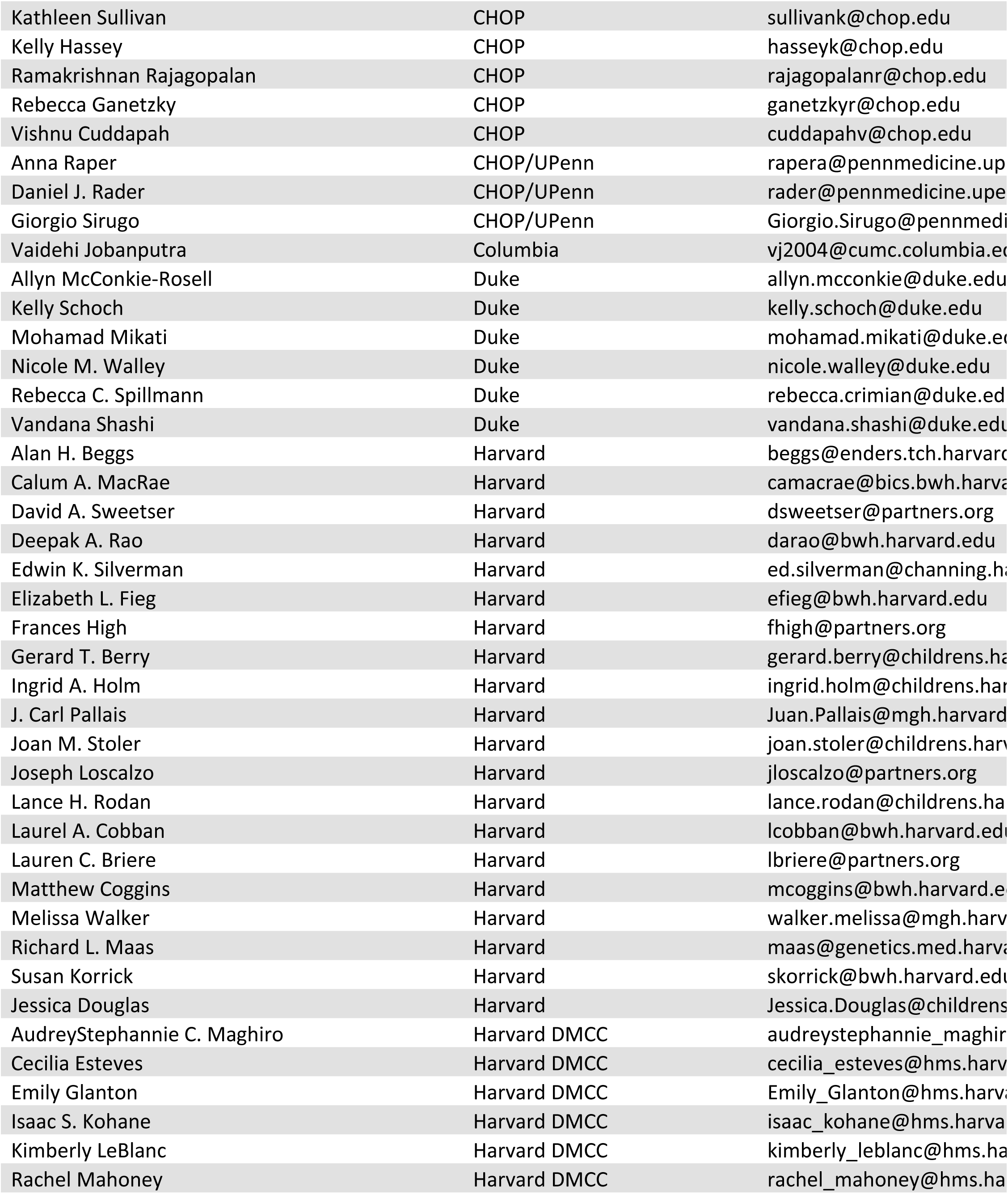
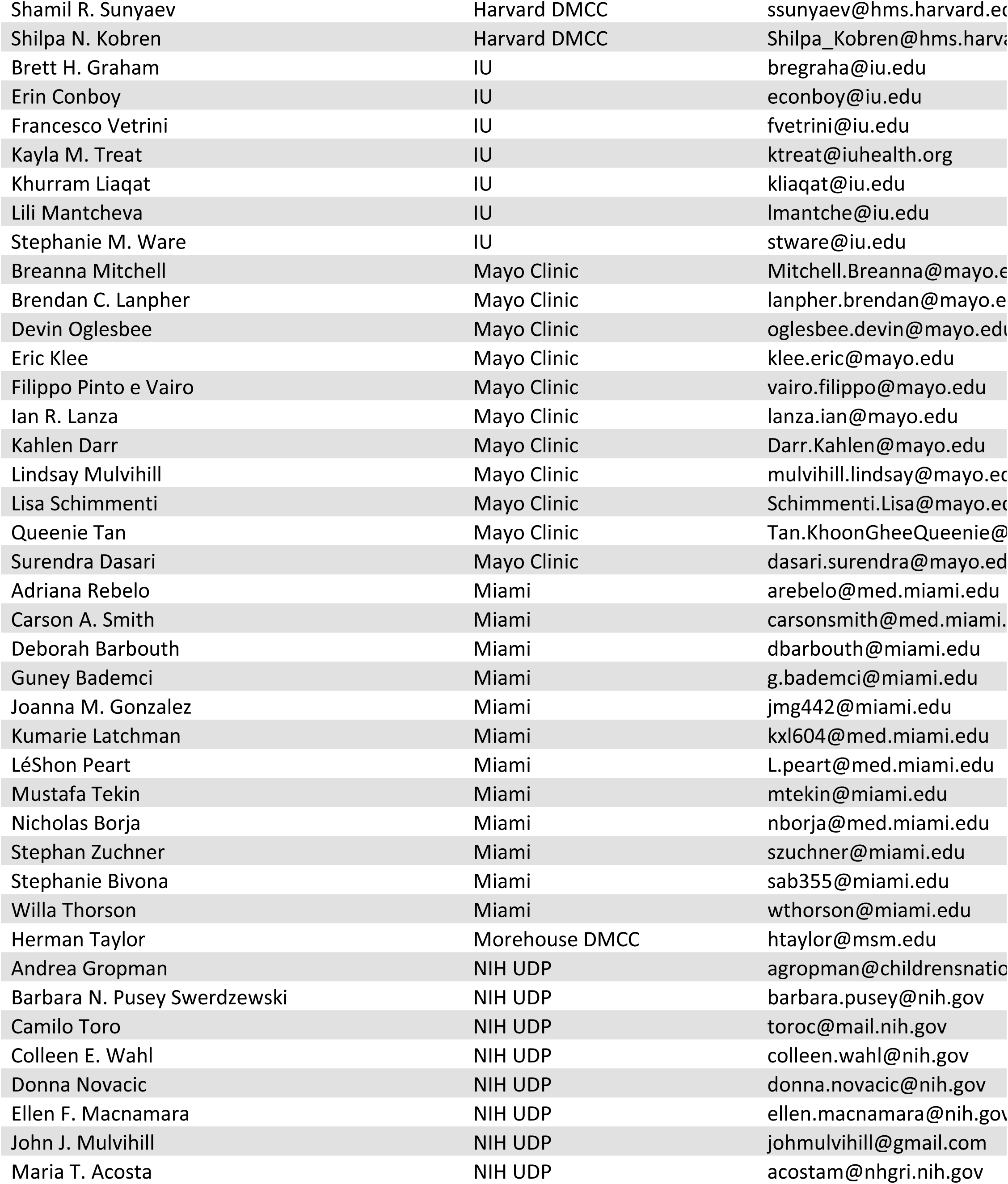
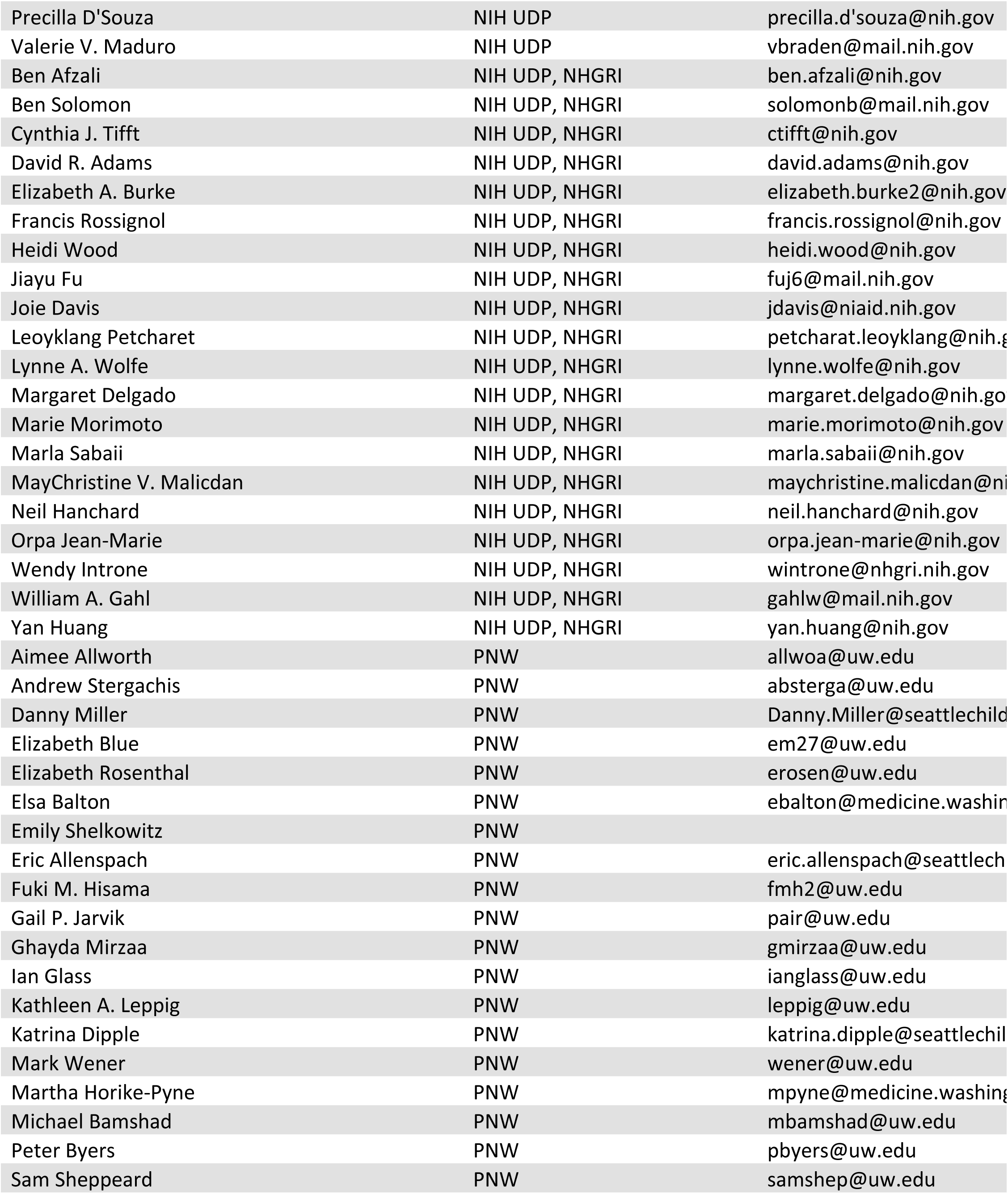
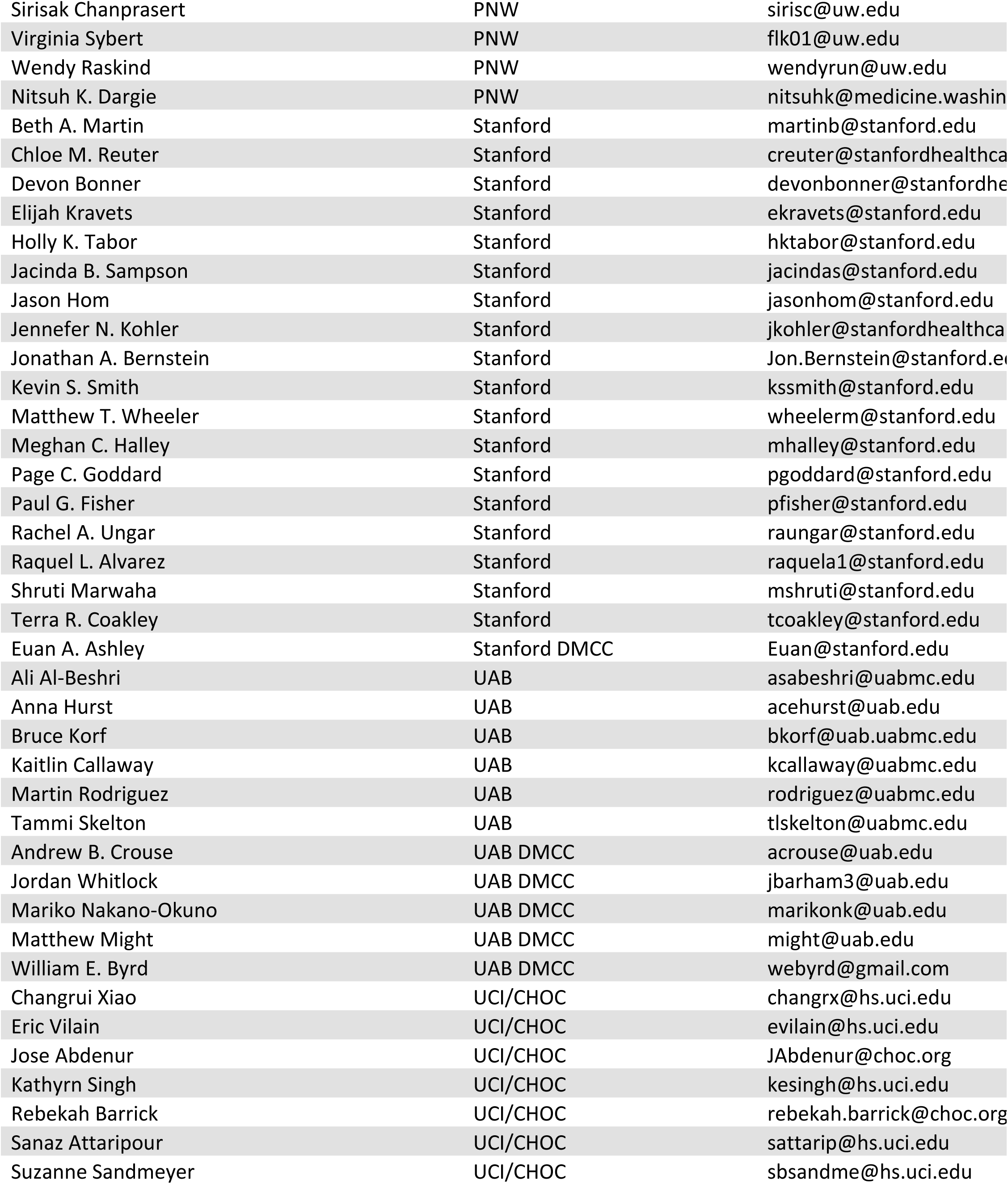
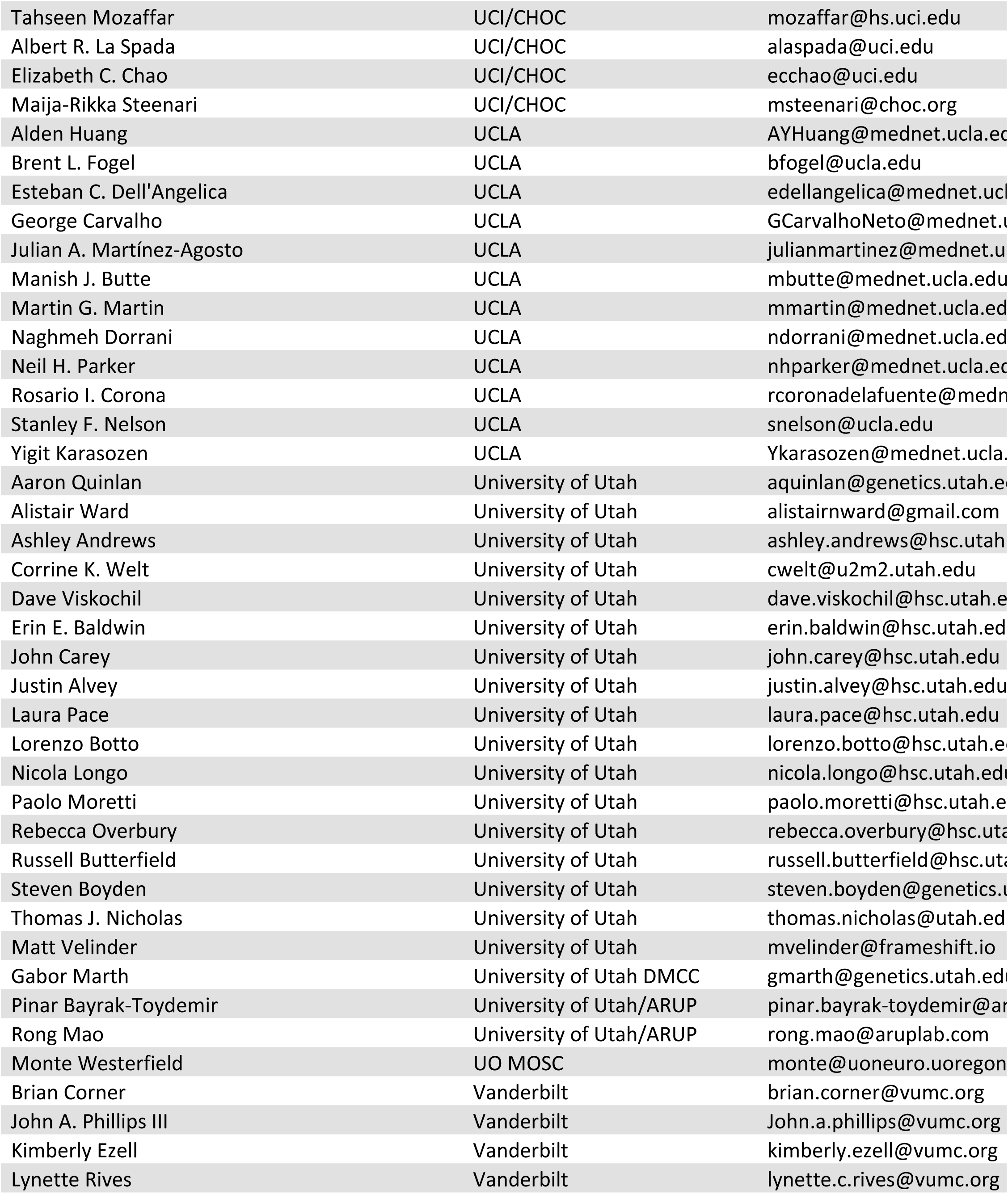
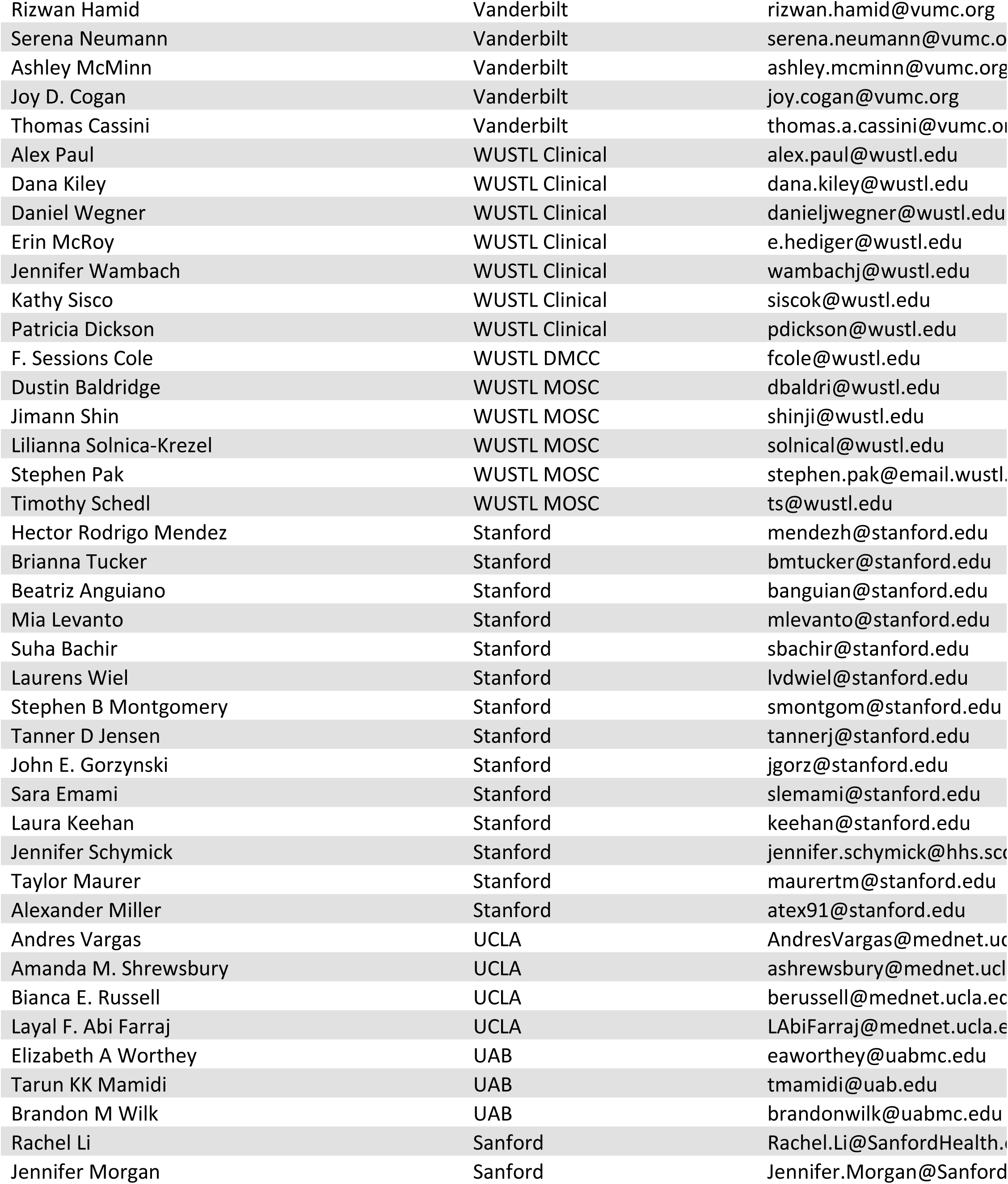
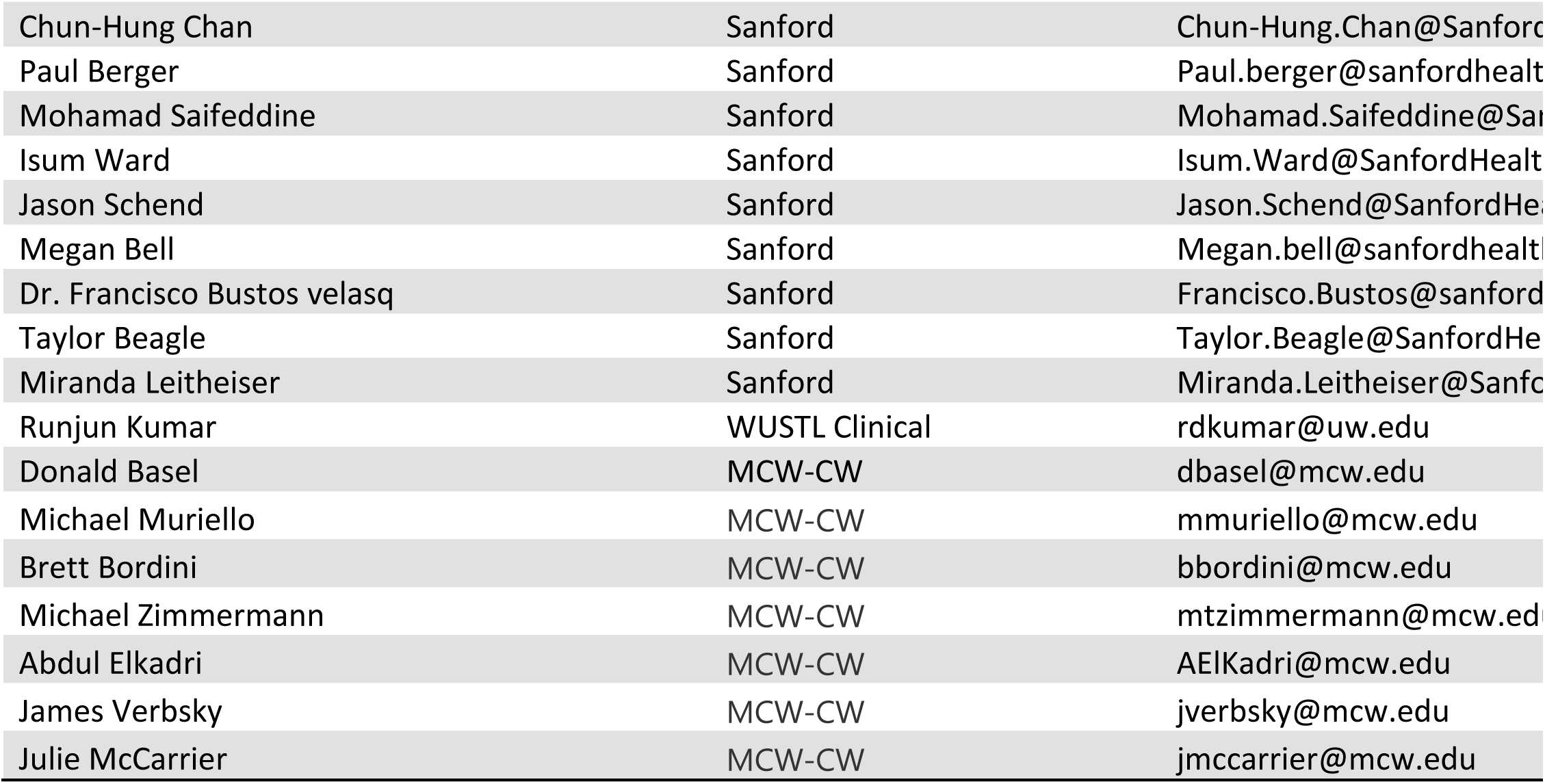

