## Supplemental Information for "VUStruct: a compute pipeline for high throughput and personalized structural biology"

### Introduction

While VUStruct has broad application to most any set of VUSs, we specifically share here the workflow we have developed to support weekly clinical cases for the Vanderbilt UDN. VUStruct allows our Personal Structural Biology (PSB) team to inform the clinic from the intersection of structural biology, genetics, and computation.

As its final automated step, the pipeline creates a patient case website of detailed calculation outputs and interactive protein visualizations. Our lead structural biologist reviews this generated website and applies her experience in computational structural biology to evaluate and curate the detailed results in broader biochemical and structural context. She particularly considers confidence and quality metrics for individual calculations and structural models. At the weekly meeting, following clinician discussions of candidate genes, she quickly summarizes the curated VUstruct outputs as a single page multicolor spreadsheet. She presents that report in context of the meeting's critical preceding discussion.

### Suggestions for Clinical Case Support

Our summary variant classifications support the discussions of the clinical team (1,2) and serve as inputs to their generation of hypotheses. In the best cases, our insights lead to testable biophysical/biochemical mechanistic explanations for pathogenicity. Uncertainties abound not only in the calculation algorithms, but also in their inputs. Given these uncertainties, the team curates raw calculation outputs through the lens of its structural biology experience. From hundred(s) of calculation details, we whittle our weekly end-product to a single page. Our manually edited case summary grid communicates, for each missense VUS (rows) and for each calculation (columns), our initial sense of whether a variant might impact function. We report our final recommendations in the right column with colors "no", "maybe", or "yes" for variants meriting further investigation.

Some calculations cannot be run due to a variety of issues ranging from lack of quality structural coverage for the variant, to insufficient available variants for PathProx spatial analyses, to variant-specific technical reasons. These missing cells are left blank in the summary. VUStruct creates this spreadsheet for each case, as an editable template. Supplemental Figure 1 is an edited summary spreadsheet from a recent case:

| Gene | Uniprot ID | variant | ddG destabilizing | PathProx 3D hotspot w/ ClinVar | PathProx 3D hotspot w/COSMIC | PPI | AlphaMissense Score | PTM | Digenic interaction by DiGePred or DIEP | Worth pursuing? | side notes |
|---|---|---|---|---|---|---|---|---|---|---|---|
| AMOT | Q4VCS5-1 | V472L | -0.5 | | 0.1 | 0.9 | 0.9 | Phosphotyrosine | | **PPI + PTM + AM** | |
| CITED4 | Q96RK1 | A106T | 9.0 | | 0.1 | | 0.07 | | | | Low confidence model |
| PAQR6 | Q6TCH4-1 | A230V | -1.1 | | 0.2 | | 0.64 | | | **ddG + AM** | |
| NLRP1 | Q9C000-1 | S38P | -1.73 - 10.41 | -0.3 | 0.0 | 0.6 | 0.06 | Phosphothreonine | | **ddG + PPI + PTM** | |
| ACADM | P11310-1 | K329E | 5.5 | 0.1 | 0.3 | 0.9 | 0.51 | | Diep w. UQCRC2, LPL, FARS | **ddg + PPI + Digenic** | variant is in oligomer interface |
| CBS | P35520-1 | R369C | 4.7 | 0.3 | 0.2 | | 0.15 | SUMOylation | | **ddG + Clinvar + PTM** | dimer interface variant |
| CDC42BPA | Q5VT25-1 | T419M | -1.8 | | 0.1 | 0.8 | 0.08 | Phosphoserine | | **ddG + PPI + PTM** | |
| CNOT1 | A5YKK6-1 | R807G | | 0.0 | 0.0 | | 0.23 | | | | |
| COX10 | Q12887-1 | C343R | 1.4 | 0.4 | 0.2 | | 0.18 | | Diep w. UQCRC2 | **ddG + Clinvar + Digenic** | |
| CR1 | P17927 | R1564Q | | | 0.0 | 0.5 | 0.09 | Phosphoserine | | | |
| DIS3L2 | Q8IYB7-1 | A83V | 7.5 | 0.3 | 0.5 | | 0.95 | | | **ddG + AM** | |
| DLL1 | O00548-1 | N300K | 4.1 | | 0.1 | | 0.93 | O-linked_glycosylation | | **ddG + AM + PTM** | |
| FARSB | Q9NSD9-1 | R98W | -2.9 | 0.5 | 0.1 | | 0.2 | | Diep w. ACADM | **ddG + Clinvar + Digenic** | |
| HYOU1 | Q9Y4L1-1 | V239I | -0.32 - 5.44 | | | | 0.08 | | | **ddG** | |
| IRF8 | Q02556 | R170G | 0.4 | 0.0 | 0.0 | 0.7 | 0.1 | | | **PPI** | |
| LPL | P06858 | N318S | -0.9 | 0.4 | 0.2 | | 0.07 | | Diep w. MSR1, ACADM | **Digenic + Clinvar** | |
| MAPK8IP3 | Q9UPT6-1 | V12M | 1.6 | 0.0 | 0.0 | 0.7 | 0.15 | | | **PPI** | |
| PIK3CB | P42338 | V199I | -1.7 | | 0.0 | | 0.08 | | | **ddG** | ras binding domain variant |
| S1PR3 | Q99500 | M365L | -1.9 | | 0.0 | | 0.07 | | | **ddG** | Low confident model |
| UQCRC2 | P22695 | T142I | 2.7 | 0.6 | 0.1 | 0.9 | 0.31 | Methylarginine | Diep w. ACADM, COX10 | **G + Clin. + PPI + PTM + Dige.** | |

| legend | |
|---|---|
| (red) | **Yes** |
| (yellow) | **Maybe** |
| (blue) | **No** |

Supplemental Figure 1. VUstruct calculation results are presented as a single case summary spreadsheet, which is manually edited from a VUStruct-generated template. The initial PSB team recommendations are shown in a "Worth Pursuing?" column and presented in context of preceding clinician discussions.

Our meeting preparations often include literature searches and give closer attention to experimentally determined structures. These insights can identify additional protein structural nuances and reveal possible variant impacts on dimerization surfaces or other interaction sites.

During the case review meeting, the PSB team lead listens closely to the case discussion and succinctly relays our findings in context. The lead doesn't report on the entire spreadsheet. Rather, she provides additional details or perspective from our results that directly support or challenge the emerging hypotheses of causal genes. The team lead may also suggest ongoing consideration of additional genes, where our calculations strongly implicate them, so long as preceding discussion has not definitively eliminated them.

Questions from the team are always welcomed and taken offline when follow-up research is needed. The PSB team is always available for additional pipeline runs, and for supplementary modeling to dig deeper into specific variants and report back on current or prior cases. Indeed, the VUstruct pipeline's integrated calculations are but a small sample of computational methods available to us.

### VUstruct Details: Calculations and their Interpretation

The pipeline-launched calculations typically output numerical values which include biophysical energy change, pathogenicity predictions, or probability of location at functional protein sites. Along with these raw numbers, the calculations deliver critically important statistical significance intervals and metrics of predictive confidence.

We now describe the specific calculations in greater detail, along with the ever-evolving interpretational guidelines that we have developed to summarize results.

### Rosetta $\Delta\Delta G_{folding}$

While proteins are tolerant of (i.e., are functional after) most single random amino acid substitutions(3,4). However, protein folding is an extraordinarily subtle phenomenon of nature. For globular protein, $\Delta G_{unfolded \rightarrow folded}$ measurements have long been understood to lie in the range [-5, -15] kCal/mol(5). Thus, the free energy of protein folding is comparable to the enthalpy of a couple of hydrogen bonds - remarkably small relative to the sum of energetics acting between all the atoms of a protein or the global loss of entropy on folding. From basic thermodynamics, it must follow that a 1.5 kCal/mol increase to $\Delta G_{folding}$ drives a 10-fold reduction in folded protein concentration and calculations have suggested that 20% of deleterious missense variants cross this low energetic barrier(6). Though the prevailing theoretical intuition around protein folding thermodynamics is arguably oversimplistic when considering more complex proteins and *in-vivo* chaperones (7), the energetic impact of a single amino acid variant can obviously be sufficient to disrupt folding. Trivial mechanistic explanations might include disruption of a critical hydrogen bond, loss of a disulfide bride, a new steric clash – any of which has the *potential* to render a protein unstable and disrupt function. Importantly, our calculations do not claim to calculate the thermodynamics of ΔG. Rather, our alchemistic hope is that the comparison of varying energetics before and after variation will be sufficient to estimate a *change* in ΔG, a $\Delta(\Delta G_{unfolded \rightarrow folded})_{variant}$.

In comparison to the subtlety of nature, our Rosetta ΔΔG Cartesian(8,9) calculations are a fast approximation. From a background total computed energy which includes hundreds or thousands of amino acid atomic interactions, (easily summing to 100,000s of kcal/mol) the Rosetta calculation attempts to determine how the new variant side chain would fit in a still-folded protein, allowing for limited rearrangement of nearby amino acids. The approximation of general protein rigidity, the treatment of only protein monomers without bound ligands, and the lack of membrane context make for a convenient and fast-running calculation with a result that we do not trust blindly. Indeed, detailed benchmarking of the computed ΔΔGs on "simple" soluble monomeric proteins found its performance only somewhat helpful in the task of engineering large libraries of protein designs, an application quite distanced from characterizing individual protein variants in individualized patients(9).

We nonetheless find the ΔΔG calculation illuminating in a *reverse* sense. When variants have a small calculated energetic impact, in the interval [-2,2] kCal/mol for our purposes, we will generally classify these as benign, depending on structural context. Outside that small range, we look more closely at 3D structures interactively, and we admit to layering subjectivity on the interpretation and classification of these results. We discount the predictive power of the calculation when it involves residues from low-confidence models, or poorly resolved experimental structure. Often, we find that the ΔΔG calculation will simply highlight amino acid changes that are predictably deleterious without any detailed calculation (ex: to or from proline or glycine, introduction of tryptophan, changes from large to small, hydrophobicity, etc.). In the end, red/yellow/green are assigned to the spreadsheet cell after some reflection. More robust analysis techniques are available if there is interest from the clinical team.

### PathProx: Pathogenic Proximity

PathProx(10,11) analyzes the placement of Clinvar(12), Gnomad(13), and random (null hypothesis) genetic variants on pipeline-selected 3D structures, and calculates whether the location of patient's VUS better clusters with the pathogenic or benign variant sets. Citation (10) offers a thorough explanation of the algorithm's mathematics and machine-learning approach. The intuitive rationale for the algorithm comes from the observation that, often, when we map variants onto amino acid side chains, we observe a higher percentage of "known pathogenic" variants clustering inside smaller spheres vs. randomly distributed variants. This technique identifies 3D hot-spots in the cartesian space of the folded protein.

For each transcript PathProx mines the Clinvar database for variants annotated as "likely pathogenic" or "pathogenic." Benign variants are harvested from the Genome Aggregation Database gnomAD and filtered for MAF > $1x10^{-5}$. Roughly, we only accept as benign those variants which are found in at least 2 gnomAD sequences, a threshold that well contrasts to the rarity of UDN patient variants.

As a control, the PathProx algorithm computes a null distribution via thousands of random distributions of both $N_{benign}$ and $N_{pathogenic}$ spheres on the then asks: "Relative to random positioning of N spheres, how do these specific N pathogenic (or benign) variants seem to cluster?". Often, we see significantly tighter clustering of the known pathogenic variants than would be expected from our random sampling. Simultaneously we see more diffuse clustering of benign variants vs. random placements. When we have both observations, and when our VUS fits more closely with the tighter clustering pathogenic set, we identify it as pathogenic, and look more closely at the 3D context of the variant sets.

PathProx cannot always make a prediction and for these cases we leave a blank cell in our summary spreadsheet. PathProx minimally requires 3 pathogenic variants, and often leave-one-out validation tells us (via low AUC), that the predictive power of the algorithm is not strong. When the calculation metrics imply confidence, we add a "no", "maybe", or "yes" to the PathProx-Clinvar column of the final spreadsheet.

As with the ΔΔG calculations, the PathProx algorithm is perhaps best validated across large protein datasets(10). Nonetheless, a strong case has been made for PathProx's power to explain disease in individual patients(11).

As with ΔΔG, the PathProx algorithm was benchmarked on protein monomers. We nonetheless run it also on multimers from Swiss-model and the PDB. We cautiously consider these multimeric calculations in our analysis – as homomeric structures introduce symmetries of variant placements that the algorithm may not be adept at scoring.

### Digenic Disease Prediction

Disease emergence through compound heterozygous variation is well understood. Individually, a Proband's disease-free parents' gene functions enough under single variation. But the combination of gene variants from each parent triggers creates the observed phenotype in the proband. Thus, compound heterozygous variant pairs are always given special attention in each week's UDN review.

Multigenic disease can analogously arise when two genes in a single pathway are damaged via variations inherited from each parent.

An increasing database of annotated digenic(14) and multigenic(15) diseases is being curated from the literature. DiGePred(16) was an initial exploration in our lab, as to whether latent digenic diseases might be predicted through machine learning strategies. Another group broadened the training inputs, while retaining a similar machine architecture, to create DIEP(17)

These algorithms are both challenged by the relatively small training data sets. DIEP has higher sensitivity, at a cost of more frequent false positive predictions. We find DiGePred is perhaps too conservative in contrast, achieving a false positive rate of 0.1% vs. 4.4% for DIEP. DiGePred is noteworthy in that it retains literature references, and outputs in an interactive JavaScript app where we can explore thresholds of confidence. We look forward to developing and integrating further developments in di- and multi-genic disease prediction.

### Protein-Protein Interaction (PPI) Site Prediction

Many proteins function through binding to other protein partners, and approximately 60% of disease-associated missense mutations have been noted to perturb PPIs(18). Amino acid variants in interaction surfaces could have negligible impact in folding energetics and still be deleterious through disruption of protein binding to usual partners. The ScanNet(19) machine learning algorithm was trained through analysis of the Dockground(20) database of 3D protein-protein interacting structures. Operationally, for each variant position covered by an Alphafold model, we ask ScanNet to predict whether the position participates in a PPI binding surface. We report any variant position to which ScanNet assigns at least 50% probability of being involved in a PPI.

### PTM

Post-translational modifications to proteins are often critical to function. However, these modifications (phosphorylation, glycosylation, and others) are typically unresolved in experimental X-ray structures, and never seen in model structures. Sequence-based prediction of post-translational modification sites has been evolving in recent decades, and we have integrated MutsiteDeep(21), the first deep-learning framework for prediction of these sites. We are particularly interested in potentially impacted PTMs in the vicinity of a VUS, and we report VUSs at sites predicted to have both at least a 50% likelihood of post-translational modification and fall within an approximate 8Å sphere around the VUS.

### CONSURF and COSMIS

Residue positions which vary infrequently across species exhibit evolutionary conservation and this is already an input to the clinic via GERP(22) and GERP++(23) scores.

As structural biologists, we are interested not only in conservation for specific amino acids in a sequence, but also in the conservation, and visualization, of the surrounding 3D protein context. ConSurf(24) colors every residue based on rates of evolution from phylogenetic tree analysis (25).

The Contact Set MISsense tolerance algorithm, COSMIS (26) was developed in our lab. In contrast to ConSurf, which quantifies evolutionary constraint based on analysis of sequence alignments across diverse species, COSMIS quantifies evolutionary constraint over more recent evolution by analyzing patterns of genetic variation across diverse human populations. It also leverages the 3D spatial context of proteins to better estimate the evolutionary constraint on protein positions of interest. Supplemental Figure 2, below, is an example depiction of COSMIS scores.

### Summary

By integrating a wide range of 3D structural calculations for each weekly UDN case, VUStruct has proven itself invaluable to the PSB team as it informs the clinic towards generation of hypotheses that explain variant pathogenicity.

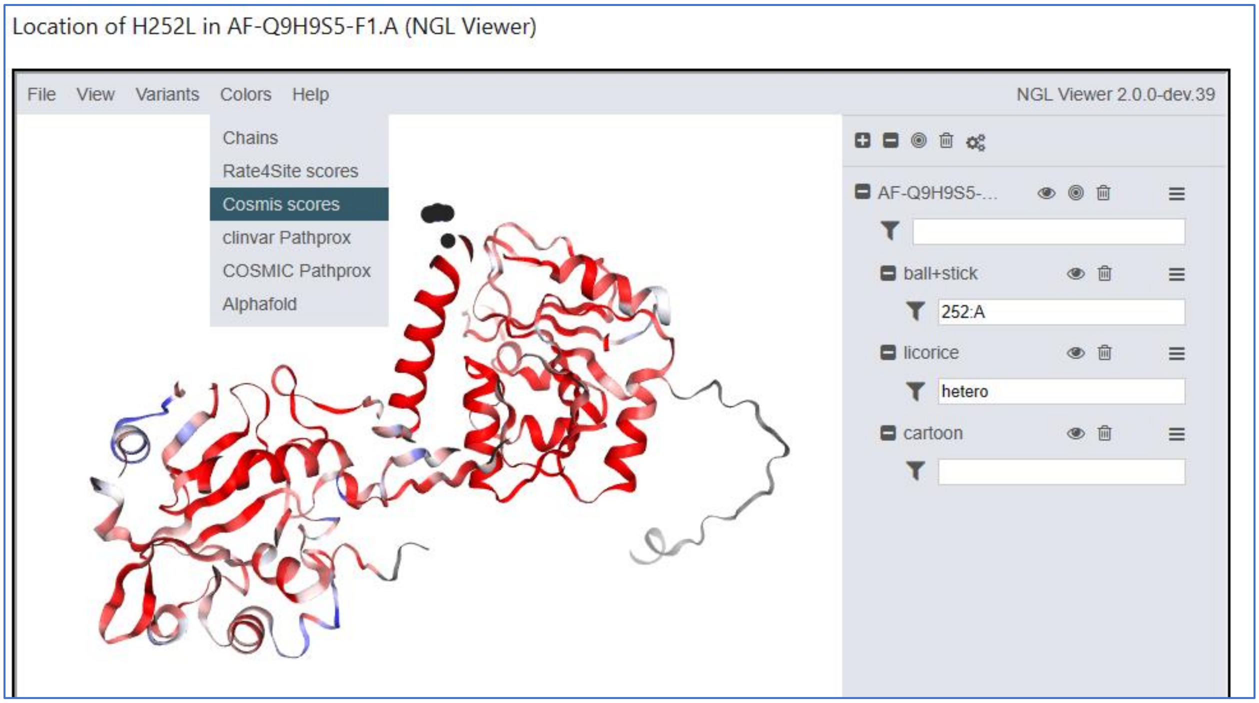


Supplemental Figure 2. COSMIS scores are depicted on the protein backbone. Red communicates high conservation. Blue tolerance to variation. This coloring scheme is also available for scores from CONSURF and PathProx. The colors in the example above reflect a general observation, that residues in highly structured protein regions tend to be more conserved across sequences, whether compared intra- or inter-species. Our pipeline additionally can rainbow-color chains in complex structures and show Alphafold pLDDTs with standard colors for those confidence metrics.
